# Three decades of increasing fish biodiversity across the north-east Atlantic and the Arctic Ocean

**DOI:** 10.1101/2022.11.03.514894

**Authors:** Cesc Gordó-Vilaseca, Fabrice Stephenson, Marta Coll, Charles Lavin, Mark John Costello

**Affiliations:** Faculty of Biosciences and Aquaculture, Nord University, Bodø 1049, Norway; National Institute of Water and Atmosphere (NIWA), Hamilton, New Zealand; School of Science, University of Waikato, New Zealand; Department of Marine Renewal Resources, Institute of Marine Science (ICM-CSIC), & Ecopath International Initiative (EII), Barcelona, Spain

**Keywords:** Climate Warming, Species Richness, Biodiversity, Arctic Ocean, Demersal fish

## Abstract

Observed range shifts of numerous species support predictions of climate change models that species will shift their distribution northwards into the Arctic and sub-Arctic seas due to ocean warming. However, how this is affecting overall species richness is unclear. Here we analyse scientific research trawl surveys from the North Sea to the Arctic Ocean collected from 1994 to 2020, including 193 fish species. We found that demersal fish species richness at the local scale has doubled in some Arctic regions, including the Barents Sea, and increased at a lower rate at adjacent regions in the last three decades, followed by an increase in species richness and turnover at a regional scale. These changes in biodiversity paralleled an increase in sea bottom temperature. Within the study area, Arctic species’ probability of occurrence generally declined over time. However, the increase of species from southern latitudes, together with an increase of some Arctic species, ultimately led to an enrichment of the Arctic and sub-Arctic marine fauna due to increasing water temperature consistent with climate change.

**Significance Statement:** Global modelling studies suggest increased species arrivals from lower latitudes and local expirations at high latitudes due to global warming. Our analysis of 20,670 standardized scientific trawl surveys with 193 fish species from the north-east Atlantic and Arctic Oceans found an increase in species richness in the region parallel to an increase in sea bottom temperature. Some Arctic species declined in probability of occurrence over time, but some increased. This, together with the increase of southern-latitude species led to an enrichment of the Arctic and sub-Arctic marine fauna attributed to climate change.

## Introduction

Climate warming constitutes one of the main faces of climate change, and is having a direct impact on species, communities and ecosystems. In the oceans, the average increase in temperature in the last 140 years has been of 1°C (1, 2). Marine ectotherms occupy most of their potential latitudinal range with regard to thermal tolerance and therefore move to higher latitudes following the displacement of their thermal niche (3, 4). However, these changes can occur at different rates across species, depending on traits such as their dispersal potential, thermal niche and capacity to exploit new resources, which may lead to changing community composition (5, 6). Understanding how these changes occur is crucial for effective conservation and management strategies. Yet, to date, consistent empirical evidence of a generalization of these shifts in the Arctic fish community is lacking.

Arctic and sub-Arctic ecosystems are among the most rapidly warming regions in the world, with some areas warming four times faster than, and seas warming twice, the global average (2, 7, 8), and their species composition may be changing accordingly (9). Until now, a doubling in species richness have been reported in some areas of the North Sea, though not in others, and increases of smaller magnitude have also been reported around North America (10–12). However, fish community analyses are mostly restricted to non-polar latitudes, and they rarely exceed the 62 °N of the north Bering’s Sea. Studies northern than this only exist in the Barents Sea, where some boreal species arrived recently (9, 13, 14) (**Table S1**). Of these, one examined distributional shifts in a fish community of 49 species from 2004 to 2017, and it reported an increase of less than 50% in species richness (14). This represents the only work reporting empirical evidence of a regional increase in fish species richness with climate change in Arctic latitudes. Thus, although an increase in species richness is predicted into the Arctic Ocean, and several model projections exist in the literature, including species extirpations (15–17), the empirical evidence is limited temporally and taxonomically, and lacks a long-term correlation with climate warming.

In the Norwegian and Barents Seas, recent warming and increased Atlantic water inflow events have been recorded, with a decline in sea-ice cover in the northern Barents Sea (18–20). As a consequence, profound effects on the geographical distribution and productivity of commercial fishing stocks are expected (21–23). In fact, species turnover was projected to increase in the area in the next decades, resulting from some local species extirpations and the arrival of warmer-water species (16, 24, 25). Here, we report three decades of field data to test these predictions.

Recent changes in species distributions in the Norwegian Sea include consistent northward expansions of the Atlantic mackerel (*Scomber scombrus*) and the European Hake (*Merluccius merluccius*), and the predicted northward expansion of bluefin-tuna (*Thunnus thynnus*), among other species (26–29). Even stronger are the changes reported in the Arctic region of the Barents Sea, where at least 11 boreal (sub-Arctic) species have been recently recorded (13, 30). Similarly, studies in the Bering Sea found localized increases in biodiversity, and suggested that areas with increased species richness and climatic stability were climate refugia (31). Several recent expansions of benthic species distributional ranges, such as that of the red king crab (*Paralithodes camtschaticus*) or the snow crab (*Chionoecetes opilio*), are rapidly altering benthic communities, and are also affecting demersal fish diets (32, 33). However, how these examples may materialise into a wider trend in the demersal fish fauna of the Arctic region due to climate change has not been investigated. Moreover, accounting for both species gains and losses needs to cover a sufficiently large region to avoid boundary effects.

Here, we test the commonness of these shifts into the Arctic across the demersal fish community of a wider latitudinal range (from 56 to 82 °N), for more species and a longer time period (27 years) than previous studies. We use three measures of biodiversity: alpha diversity (average local species richness), beta diversity and its component turnover (excluding species richness effect), and gamma diversity (total regional species richness) (34–36).

## Results

### Alpha diversity

A total of 193 demersal fish species were recorded between 1994 - 2020 across the whole study area (**Fig. 1, Table S2**). The generalized additive models (GAM) used to account for the differences in sampling effort across time, predicted a 68 % increase in the average number of species per trawl (alpha diversity), from 8.1 species/trawl in 1994 (95 % CI 7.9, 8.3), to 13.6 species/trawl in 2020 (95% CI 13.3, 13.9) (**Fig. 2, Table 1**). The increase in species richness was correlated with changes in sea bottom temperature (Pearson r = 0.59, p< 0.05, **Supplementary Results**). Increasing species richness was also found for the adjacent areas in the North Sea and around Svalbard though no significant correlation was detected with SBT at those regions (**Table 1, Fig. S1**).

**Table 1.** Average species richness per trawl (alpha diversity) in the first and last year of sampling predicted from a GAM with Year and Latitude as smooth predictors, and swept area as a logarithmic predictor.

**Figure 1.**
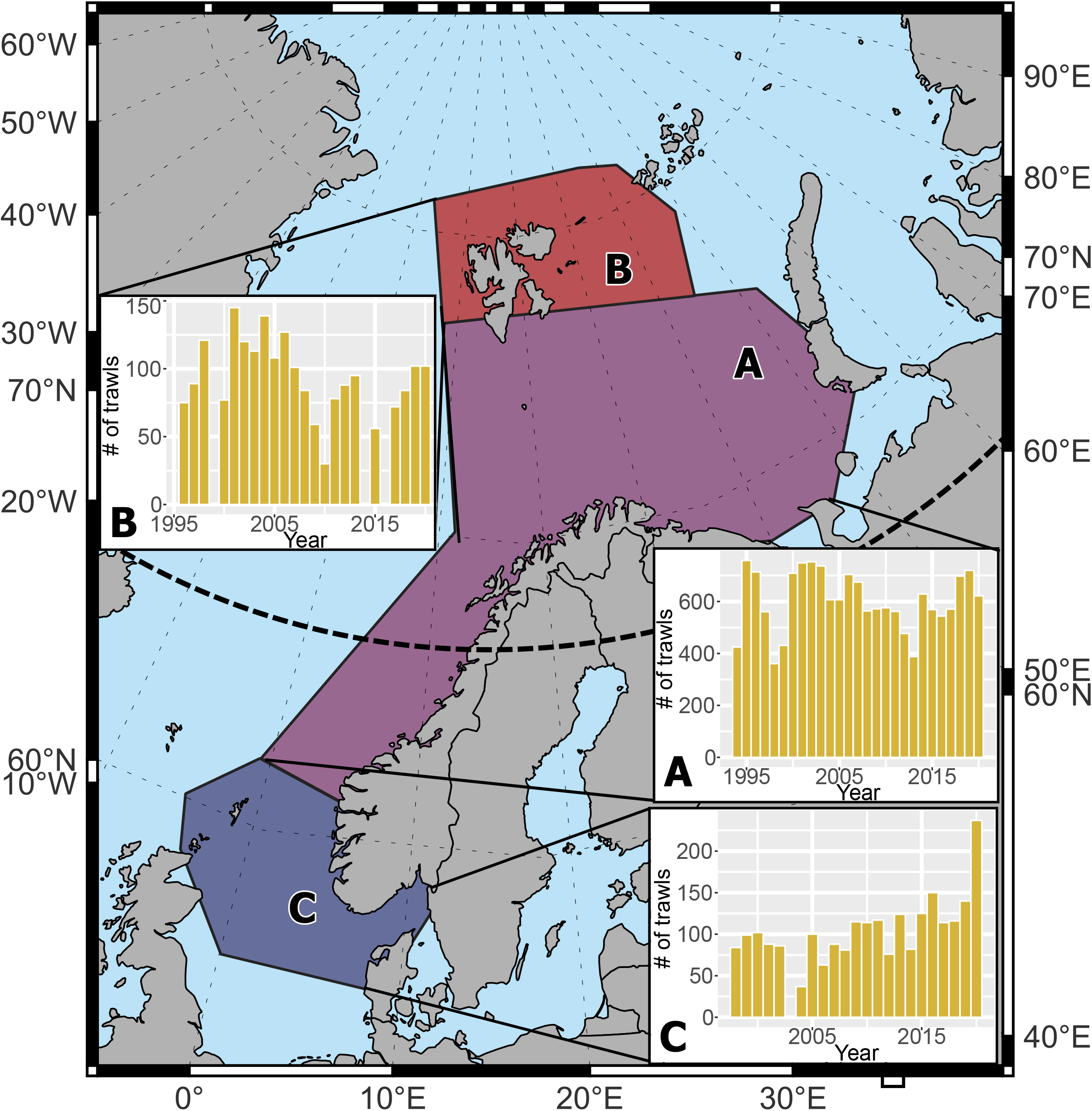
Study area and histograms of the temporal distribution of the number of trawls in the present study. A: Norwegian & Barents Sea; B: Svalbard; C: North Sea. Dashed lines are latitude and longitude, and thick black dashed line is the polar circle (66 °N).

**Figure 2.**
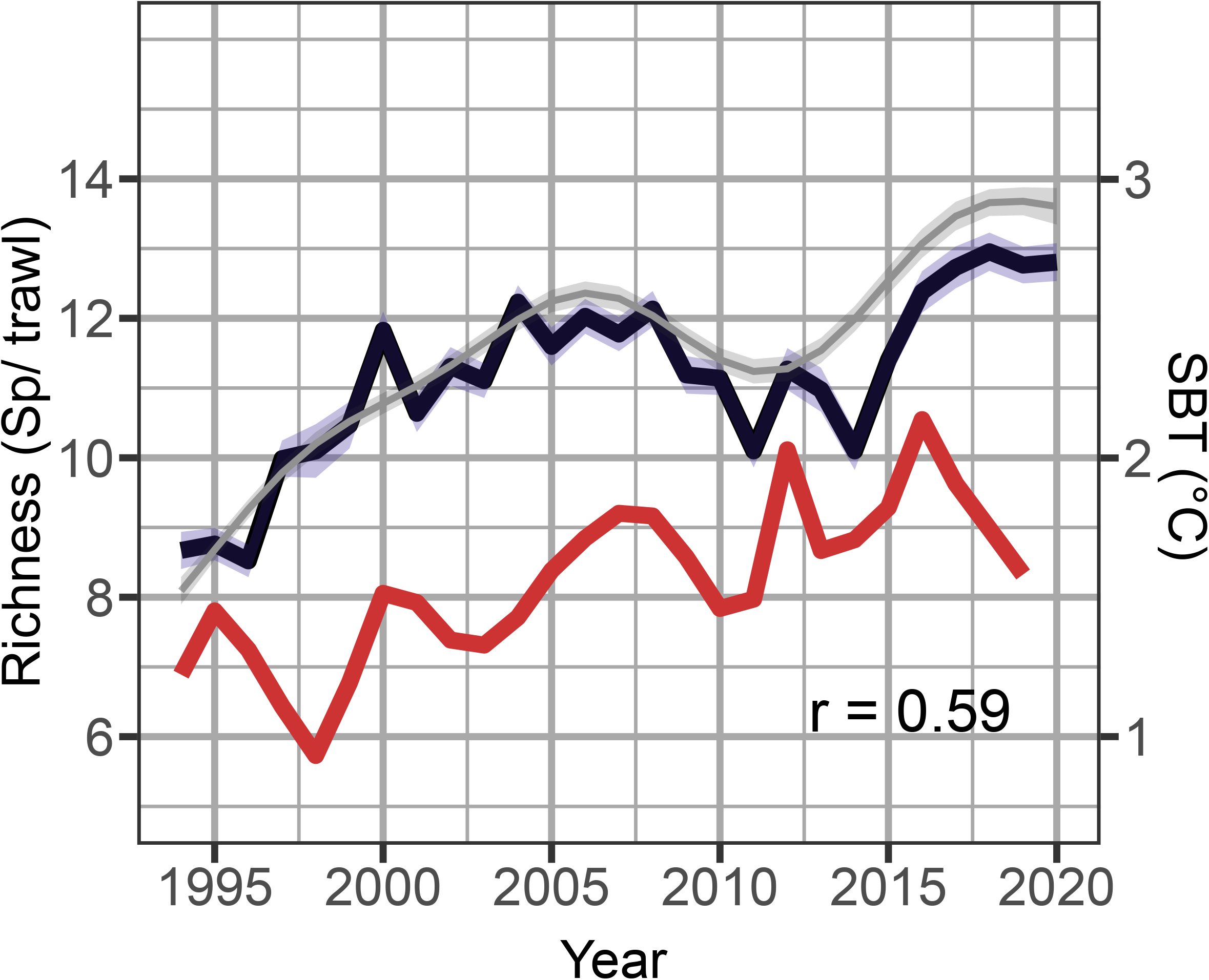
Average number of species per trawl (alpha diversity) across the Norwegian - Barents Sea. Black line represents mean species richness per trawl with 95% CI in blue. Grey smoothed line and light grey 95 %CI represents the marginal effect of Year for constant sampling effort from a GAM model using Year and swept area as an offset (**Table 1**). Red line indicates changes in mean Sea Bottom Temperature across the whole area (correlation with mean alpha diversity r = 0.59).

The boosted regression tree model (BRT) predicted local changes from 0 to 125 % in local species richness from 1994 to 2019 (correlation with independent data r = 0.63, DV explained = 38 %, p< 0.05). Among the 17 explanatory variables included in the BRT model, “Depth” was the most relevant, followed by Year and “Bottom temperature” (**Fig. S2**). The model predicted an increase in species richness at higher latitudes, especially high in the Arctic region of the Barents Sea, where species richness more than doubled in some regions, during the study period. (**Fig. 3, Fig. S3**).

**Figure 3.**
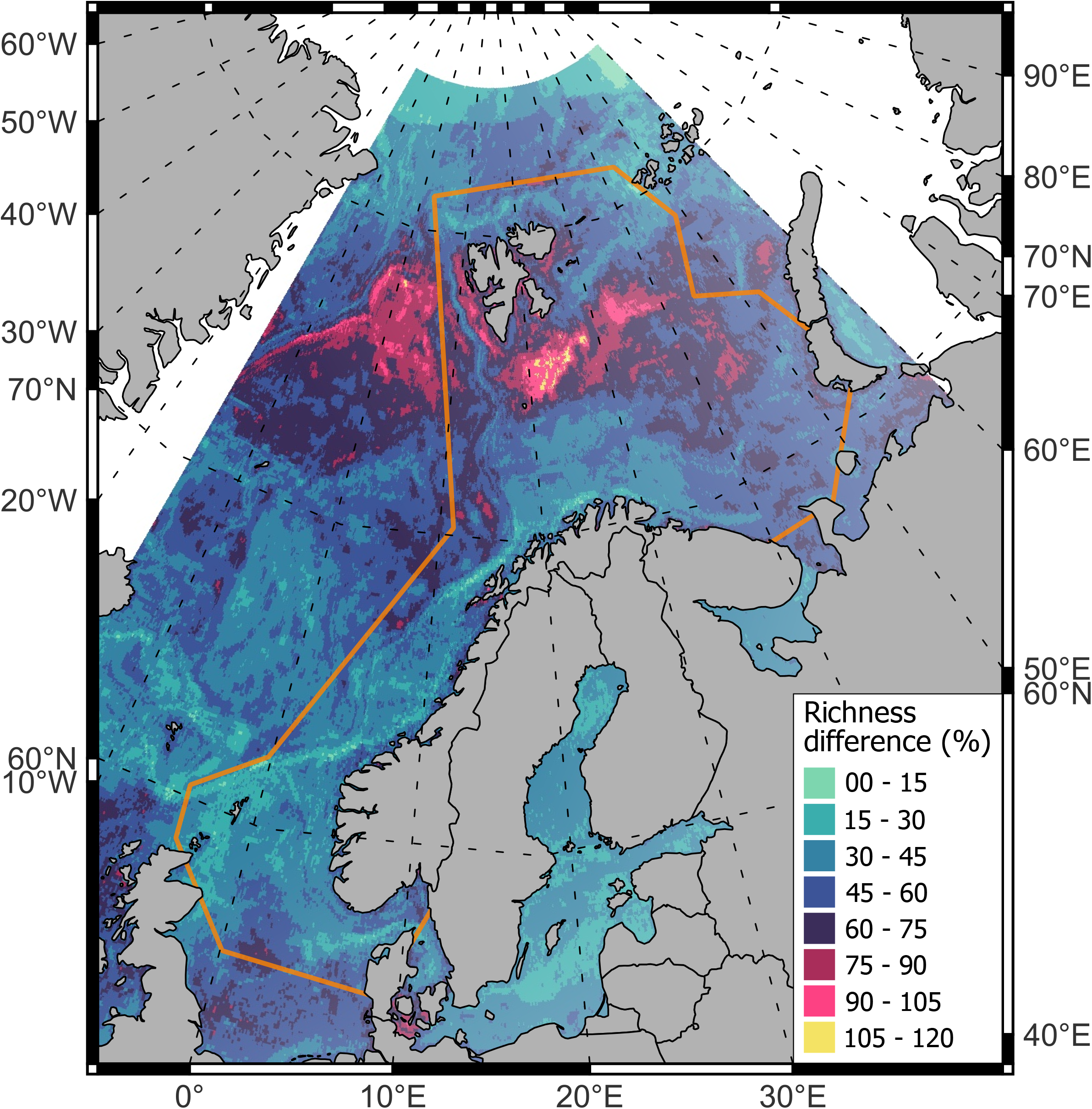
Difference between mean species richness from 1994-1996 and 2017-2019 expressed as percentage of change. The orange polygon is the study area boundary. Dashed lines are latitude and longitude.

### Gamma diversity

The overall species richness in the main study area (gamma diversity), increased 45% during the study period, from 65 species in 1994 to 94 species in 2020, following a similar temporal pattern to alpha diversity (**Fig. 4, Table 2, Fig. S4**). Very similar increases were reported in adjacent areas (47 % in the North Sea, 45 % in Svalbard, **Table 2, Fig. S4, Fig. S4**). GAM models for total species richness at each region, with Year and both annual mean trawl-swept area and total mean swept area only selected Year as relevant explanatory variables. All additional estimators, including Chao index, incidence based (ICE), and first and second order jackknife estimators, calculated across the study area, detected an increase in gamma diversity across time, of similar magnitude (**Fig. S5**).

**Table 2.** Total species richness predicted from GAM with Year as the only explanatory variable at each region (gamma diversity) in the first and last years of data, at minimum common annual trawls.

**Figure 4.**
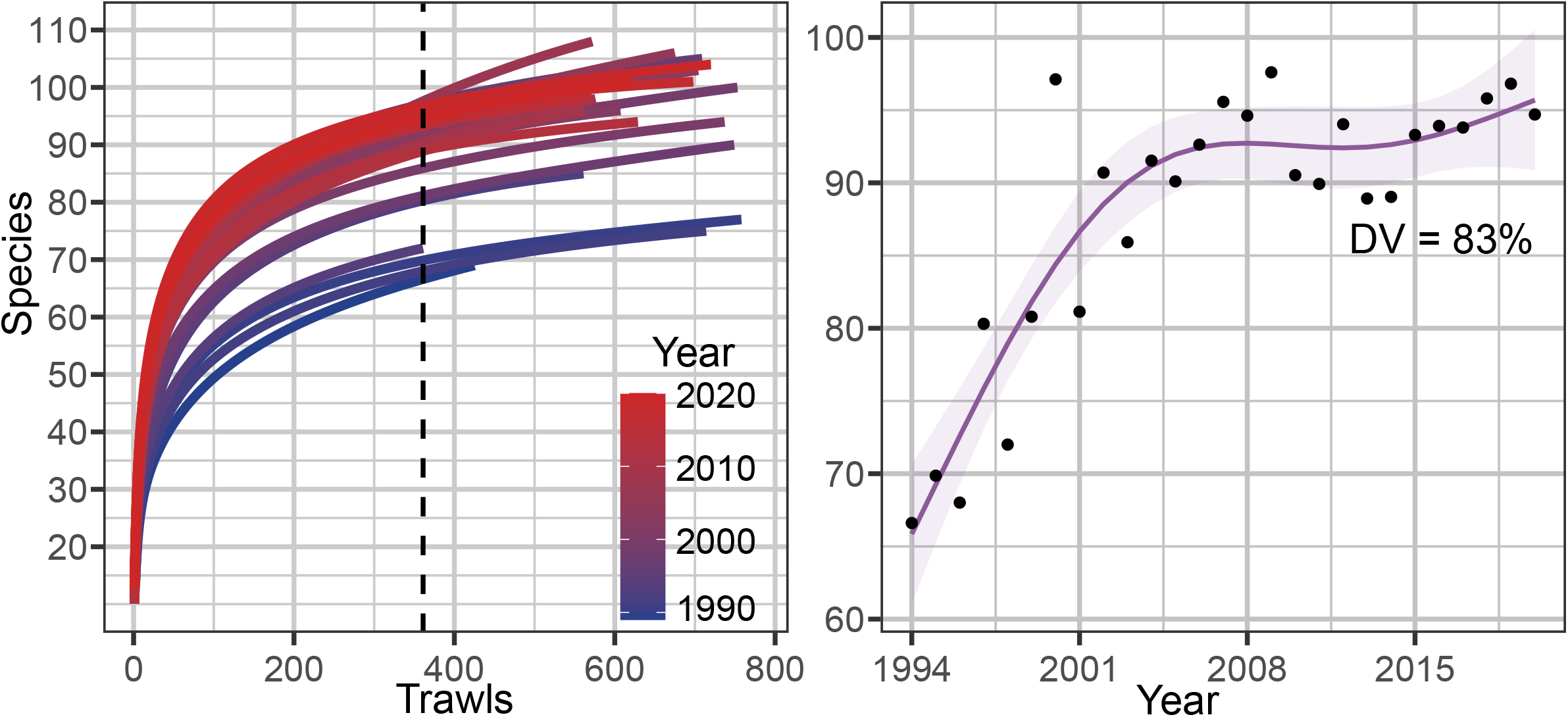
Left: Annual species accumulation (rarefaction) curves in the main region Norwegian-Barents Sea. Dashed line marks the minimum common number of sites. Right: Annual mean accumulated species richness within the minimum common number of trawls.

### Beta diversity

Pairwise mean total beta diversity increased from 1994 to 2002 in the main area, slightly declined until 2012, and slightly increased again until 2020, with an overall non-significant increase of 4 % (95 CI -3, 9) (**Fig. 5**). The turnover contribution to beta diversity, which is not affected by species richness, increased until 2008, declined until 2014, and increased afterwards, with an overall increase of 16 % (95 CI 6, 27) (**Fig.5, Table 3**). Similarly, total beta diversity in the adjacent region of Svalbard increased until 2005 and declined afterwards. However, the turnover increased linearly across time in this region.

**Table 3.** Mean total beta diversity and turnover predicted from GAM at first and last year of data at each region, at minimum common annual sites.

**Figure 5.**
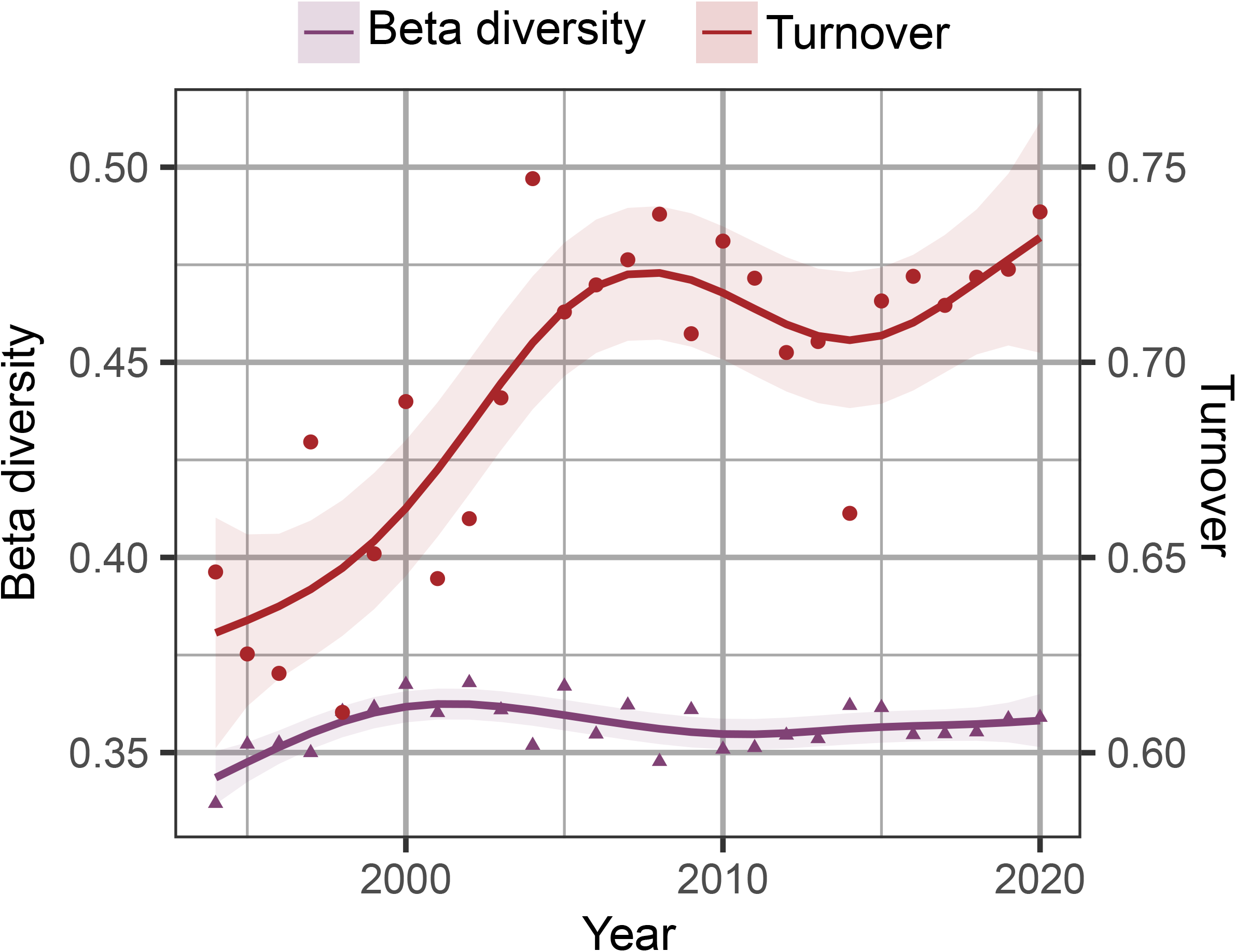
Annual mean beta diversity and turnover (richness independent), red line) in the main region across time, calculated from mean pairwise Jaccard dissimilarity index.

In the North Sea, both the total beta diversity and turnover declined with time (**Table 3, Fig. S6**). The GAM models for temporal change in beta diversity and turnover, did not select trawl-swept area as a relevant explanatory for the main study area or the adjacent regions. Thus, when the increasing species richness is accounted for, species turnover beta diversity, as well as alpha and gamma diversity increased over the years.

### Species’ trends

From the 193 species in our study, 99 species showed a significant temporal trend (increasing or decreasing) in their probability of occurrence with time, in at least one of the studied areas (**Fig. 6**). Of these, 71 species showed only positive trends, and 23 species showed only negative trends in at least one studied area, while 5 species showed positive and negative trends depending on the study area (**Fig. 6**). Thus, while no trend was detected for about half of the species, 76 % of species showing a significant consistent trend, increased. The number of species increasing was consistently higher than the number of species decreasing over time, across all regions. Species declining mostly inhabited high mean latitudes, and linear regression identified a negative effect of mean latitude on the temporal change in species probability of occurrence in the main study area, and around Svalbard (**Fig. 6**). Among the 67 species only found in the main region and/or Svalbard (Arctic region), 18 species showed a significant change in their probability of occurrence with time, 6 increased and 12 declined.

**Figure 6.**
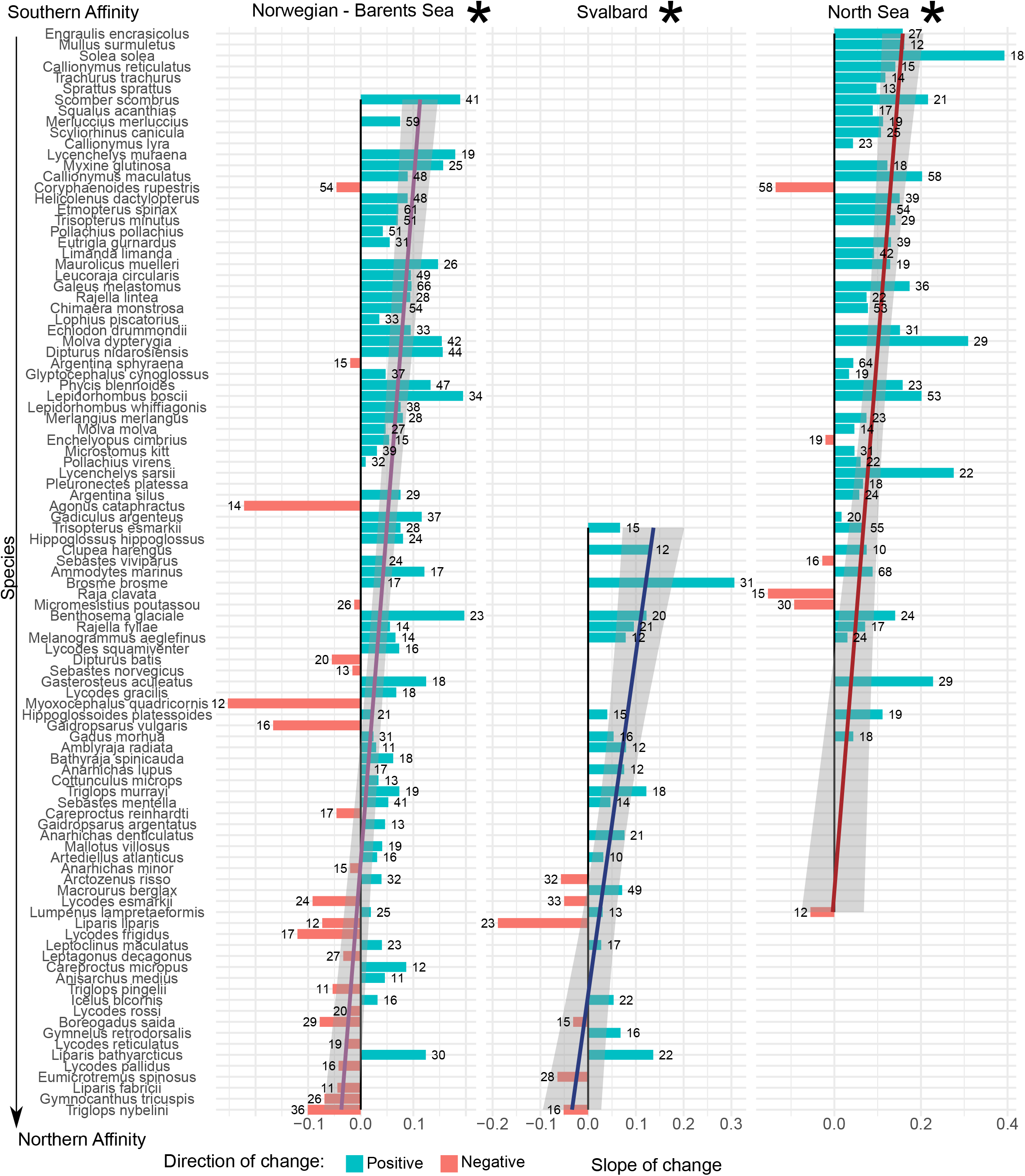
Change in the probability of species’ occurrence with time. Length of the bars correspond to the Slope of the effect of Year on species probability of occurrence, using a GAM model including swept area, Latitude and Depth as smooth predictors, and Year as a linear predictor. Species are ordered by their mean latitude of occurrence during the study period. Linear regression on the effect of Year with mean latitude was conducted and plotted, and asterisks indicate significance (p < 0.05).

## Discussion

Following global trends in warming, sea bottom temperature data suggests an increase of 0.3 °C per decade in our study area from 1993 to 2019, and this increase was several times higher in some regions of the Arctic Barents Sea (8). Here we show that this has been accompanied by a significant increase of the demersal marine fish species richness during the last three decades, in alpha, gamma, and turnover beta diversity. Considering previous studies focusing on smaller areas and fewer species (14, 31), this study indicates an ecologically significant increase in species richness in the Atlantic side of the Arctic in concert with climate warming. Consequently, species are expanding their range poleward, directly increasing the local species richness, and probably increasing functional diversity in the Barents Sea, in line with other results in near-Arctic regions (10), and with predicted poleward shifts in the North and Arctic Seas (15). This increase in biodiversity was concentrated in two different periods, interrupted by a decline period in alpha, beta and gamma diversity between 2007-2014, as suggested by Ellingsen et al. (2020).

Even though there have been studies showing consistent changes in biodiversity patterns and beta diversity at a global scale, whether a systematic loss of local (alpha) diversity has or will take place remains uncertain (38–40). Several species are changing their distributional range, mostly expanding poleward, which may eventually lead to redistributions of global biodiversity and local increases in biodiversity (10, 38). In an analysis of 50,000 marine species, Chaudhary et al. (2021) found that thousands of species had shifted from equatorial to mid latitudes, and using independent data not available to that study we confirm that this shift has continued into higher northern latitudes, leading to two times more demersal fish species in the study area. Increases in alpha diversity were already reported in some regions of the Arctic Ocean, albeit smaller areas and/or shorter time periods (14, 31). With this study, we show that the increase in species richness is not localized, but widespread across the region and concomitant with the increase in bottom temperature.

We identified sea bottom temperature as the most relevant temporal environmental variable in our model, and the third variable after Depth and Year. The fact that Year remains an important variable, and that the model explains 36% of the overall deviance in species richness, suggests that other processes and variables are needed to fully explain the changes in species richness across the study area during the study period. Still, the increase in species richness found on the continental shelf enriches the food web of the area, where most socio-economic activities occur, compared to the open ocean. Variations in food-web structure have been already reported in the Barents Sea, across environmental gradients, with temperature as one driver of these changes (30, 41). Increased competition has been reported between some gadoids, which may have a detrimental effect on the abundance of polar cod (*Boreogadus saida*) as the abundance of boreal species increases (42). Polar cod is a key species in the Arctic food web (43) and its decline could affect the subsistence fisheries of native communities that harvest higher trophic level species (44). Ultimately, changes in biodiversity will interact with other multiple effects of climate warming, as changes in species distributions and food-web rewiring will likely affect ocean productivity and fisheries (25, 45–47).

Most studies on species richness consider only gamma (total) or alpha (average) diversity. When beta diversity is considered it has most commonly been measured as Jaccard’s coefficient which is not independent of species richness (48). Here, we find that while both alpha and gamma diversity are significantly increasing, so is species-richness independent species turnover, but not total beta diversity (**Fig. 5**). Thus, the spatial heterogeneity of biodiversity is increasing as well as biodiversity overall. However, the difference between alpha and gamma diversity changes in our study is likely not large enough to reflect a change in overall beta diversity.

Overall the study area, we found that 76% of the species showing a consistent significant change in probability of occurrence increased with time. However, when restricted to species absent from the North Sea, 67 % of them declined with time. When species’ mean latitude was used as an indicator of the Arctic affinity of each species, it significantly explained the effect of year in species probability of occurrence in the Norwegian and Barents Seas, and also around Svalbard. Although this a clear indicator of a decline of several Arctic species, not all species occupying high mean latitudes showed a decline in probability of occurrence with time. Some high-latitude species increased substantially, perhaps because they benefited from changes in food-web interactions. This also suggests a partial coexistence between boreal and Arctic species that, together with the increase of lower-latitude species, leads to a consistent increase in biodiversity, in line with previous results on fishes and crustaceans (42, 49), but not with a previous hypothesis of a synchronous species extirpation in the western Barents and Norwegian Seas (16, 24). If this trend is maintained in the following decades, we may observe enriched communities in the Arctic, which may be sustained by the projected increase in net primary productivity of up to 50% (25, 50). The wider effects of changing fish biodiversity on marine food webs remains to be investigated.

## Materials and Methods

### Study area

The database here analyzed comprises research trawl surveys restricted to the continental shelf and slope from the north of the North Sea into the Arctic Ocean, mostly from 56 ° N to 81 ° N Latitude, and from 2 °W to 50 °E Longitude (51). The whole area comprises a marked temperature gradient, with average bottom water temperatures over 8 °C in the North Sea to -1 °C in the northern region of the Barents Sea. Because of data temporal and spatial distribution, we considered one main study area, with the longest time period (Norwegian & Barents Sea) and two adjacent areas (Svalbard and North Sea), which contain a shorter temporal period (**Fig. 1**).

The database initially contained 60,355 research surveys, from 1980 to 2020. We excluded data which were associated with broken gear, had incomplete metadata (data without reporting depth, or coordinates, or type of gear), and were questionable (e.g., shrimp trawl opening of several km). We restricted the analysis to shrimp trawling data using 20 mm mesh size, a maximum of 5 km trawling distance and 60 m of trawl opening, from 30 m to 700 m depth. We only included fish species (classes Actinopterygii, Elasmobranchii, Holocephali, Myxini and Petromyzonti). Invertebrates were not included in the study. After data standardization and selection, 20,670 surveys from 1994 to 2020 remained.

Depending on the data spatial coverage over time, and based on the biogeographical realms that have been described in the area (52), we divided the study area into a main area containing most of the data and the longest temporal coverage, the Norwegian-Barents Sea, and two adjacent areas with more limited data: the area around Svalbard and the North Sea (**Fig**. 1). Although the latitudinal and longitudinal distribution of trawls across time was relatively homogeneous at each region, the swept area of each trawl presented a weak negative trend with time for the Norwegian-Barents Sea and Svalbard regions, which was accounted for within the statistical modelling (**Fig. S7, Fig. S8**).

### Environmental variables

Spatially explicit environmental co-variates were collated from three sources: Copernicus Marine Service database, Bio-Oracle and MARSPEC database (53–55). Some of the environmental variables were available as annual estimates (e.g., sea surface temperature) whereas others were available as long-term averages (e.g., bottom nitrate concentration) (**Table S3**).

Variables from the Copernicus Marine Service were available for each year from 1993 to 2019. These include estimates of sea surface temperature (SST), sea bottom temperature (SBT), sea surface salinity (SSS), ice concentration and sea surface currents (northward and eastward components), and were obtained from the “GLOBAL_REANALYSIS_PHY_001_031” dataset (55) (**Table S3**). Because these layers were only available from 1993 to 2019, we limited the analyses to this period. Long-term averages (2000 – 2014) of nitrate, iron, dissolved oxygen, surface and bottom productivity, and temperature range were obtained from Bio-Oracle. Bathymetry was obtained from the MARSPEC database (53, 54). Variables from Bio-Oracle and MARSPEC were downloaded using the “sdmpredictors” package from R software (56).

A distance to coast layer was created using the “raster” package, also in R (57). All the environmental layer resolutions were matched by downscaling to the lowest original resolution (0.083 ° Longitude x 0.083 ° Latitude). Variables from Year, Latitude, Longitude and Annual trawls were also manually created. The shapefiles for the ocean and land were obtained from the Marine Regions database (58) and maps were created using QGIS (59).

The change in SBT with time was calculated as the annual mean across each study area (**Fig. 2, Supplementary information Methods**). SBT thus reflects the likely average temperature individual fish may have experienced over time rather than temperature when sampled.

### Biodiversity measures

Species richness is typically measured as alpha (local) and gamma (regional) diversity, while the extent of differentiation of communities along habitat (spatial and/or temporal) gradients, is beta diversity (35, 60). The most commonly used measures of beta diversity, such as the Jaccard index, are influenced by species richness, which should be accounted for where species richness significantly varies. We thus report both total beta diversity and a species-richness independent index here called turnover, following (34). The mathematical difference between beta diversity and turnover is called nestedness, and arises where sites with less species are a subset of species from neighbouring sites with more species. Thus, we report four measures of diversity: alpha, gamma, total beta, and turnover.

### Alpha diversity

We assessed alpha diversity in the study area as the annual mean species richness per trawl (species/trawl) using presence data only. Generalized Additive Models (GAMs) were used to explore the changes in species richness with time, correcting for the different swept area by adding it as a logistic explanatory variable. The model construction was done using the package “mgcv” within the R software (61, 62).

We used boosted regression trees (BRT) to explore the drivers of these changes, and to make spatial projections of the species richness in the study area. A threshold of 0.9 correlation coefficient was selected due to the robustness of the procedure to collinearity (63) (**Supplementary information Methods**). We restricted the modelling period to the years 1993 – 2019 due to the restricted availability of contemporaneous environmental data in the Copernicus Marine repository. The data were randomly divided for model assessment: 75% was used for calibration, and 25% for model validation (**Supplementary information Methods**).

To study the effect of each environmental variable on species richness, we built partial dependence plots using the R package “devEMF” and basic R from the BRT model. These showed the mean species richness predicted across each environmental variable gradient, while all the other variables were kept at their means (61, 64).

To explore the spatial changes in alpha diversity over time, we statistically predicted the geographic distribution of species richness in each year from 2017-2019 and 1994-1996 from the BRT model, and calculated the difference between the mean of each period. We used the “predict” function of the “dismo” package, and the package “raster” from R (57, 61, 63, 65, 66). Maps were created with QGIS(59).

### Beta diversity

Beta diversity and its turnover component were calculated using the mean pairwise Jaccard dissimilarity index (ß_J_) from presence-only data, which can be divided into two components: species replacement (Repl.) and nestedness (Nest.) (34, 36, 67):

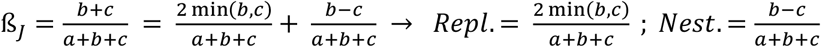

Where *a* refers to the number of species present in both sites, and *b* and *c* are the species present only in either site. Pairwise means each trawl is compared with every other trawl in a year and the average dissimilarity calculated.

Here we report total beta diversity (ß_J_) and the relative contribution of species replacement, to beta diversity, which we refer to as turnover:

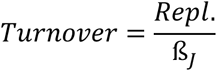

To account for differing sample sizes between years we conducted a bootstrapping process. That is, we randomly selected a subsample of the smallest sample size over years, calculated its total beta diversity and turnover, and repeated this process 200 times for each year (34). We report the mean of these bootstrapping calculations in **Figure 5** and **Figure S6** (34). All calculations were carried out using the functions beta.div.comp, from the adespatial package (68).

To explore the trend of these changes with time and to account for the negative trend of swept area with time (**Fig. S3**), we fitted a GAM model with Year and annual mean swept area as explanatory variables to the annual mean pairwise total beta diversity and turnover.

### Gamma diversity

To study changes in gamma diversity, we constructed species accumulation curves (SAC) for each year across our study regions (69, 70). To eliminate the bias that a selected order of the locations may have on the overall SAC, we used the function “specaccum”, in the “vegan” package, set the parameter method = “random” to randomize the order of site addition, and permutated the process 200 times (70, 71). We report the mean species richness per trawl of all random permutations. We then assessed the change in gamma diversity with time, standardized to the minimum common number of sites, fitting a GAM model with Year and annual swept area as the explanatory variables.

To test if the results were robust to alternative measures of richness, we used the package “SpadeR” to calculate nine different species richness indices with time, each of them with different statistical assumptions, including Chao indices, incidence based indices, and first and second order jackknife estimators (72, 73).

### Individual species contributions

To study which species drove the changes in biodiversity, we fitted generalized additive models to the presence data of each species with smooth parameters for depth, swept area and latitude, and fixed effect of Year, using a binomial distribution. Each species abundance-weighted mean latitude was calculated from the complete area to arrange the species by their Arctic-affinity, and simple linear regression was used to assess the effect of mean latitude on the temporal change in species probability of occurrence.

## Acknowledgements

C.G.V would like to thank Eric Molina for his interest in the work and helpful discussion. MC would like to acknowledge partial funding from the Spanish National Project ProOceans (PID2020-118097RB-I00) and institutional support of the ‘Severo Ochoa Centre of Excellence’ accreditation (CEX2019-000928-S). FS would like to acknowledge partial funding from the National Institute of Water and Atmospheric Research (NIWA) Coasts & Oceans Research Programme 5 (SCI 2020/21).

