## Supplementary material for "Three decades of increasing fish biodiversity across the north-east Atlantic and the Arctic Ocean"

**This PDF file includes:**

Supplementary text

Figures S1 to S8

Tables S1 to S3

**Supplementary Information Text**

**Results**

**SBT model.** A linear regression with time, longitude and latitude as the explanatory variables for the changes in SBT from 1993 to 2019 in the main study area explained 20% of the deviance and suggested a rate of warming of 0.03 °C/yr.

**Methods**

**Boosted Regression Trees** Environmental variables correlated higher than pearson = 0.9 were eliminated to deal with extreme collinearity (BRT are highly robust to collinearity),which was not met between any variables. Additionally, variables that did not improve the model were also eliminated to keep the model as simple as possible without compromising model quality (**Table S3**) (meaning not improving total explained deviance and individually accounting for < 4 % of deviance explained) To improve the model calibration and reduce computational requirements, we conducted a bootstrapping process: we fitted a model to 75% of the trawls, and predicted each year’s species richness distribution with it. Then, we repeated this process 50 times and reported the average of the 50 models per year. Each time, the data used for calibration was a random selection with replacement, of the size of the whole calibration dataset (75% of trawls). Each model was limited to 20,000 trees, and tree complexity was set to 3 to allow for interactions between the explanatory variables. The learning rate was adjusted to 0.01, bag fraction to 0.3, and other parameters were set to their defaults. Model validation was conducted through Pearson correlation with the remaining 25% of the dataset that was not used for model calibration.

Richness was projected annually for each of the 50 models from bootstrap, using each year’s environmental layers. We additionally created a “swept_area” raster file using the unique value of 28 km^2^ across the whole area, which maximized the species richness in the BRT partial dependence plots and corresponded tot the maximum swept area within the dataset. Annual mean projections were obtained from the 50 projected richness rasters.

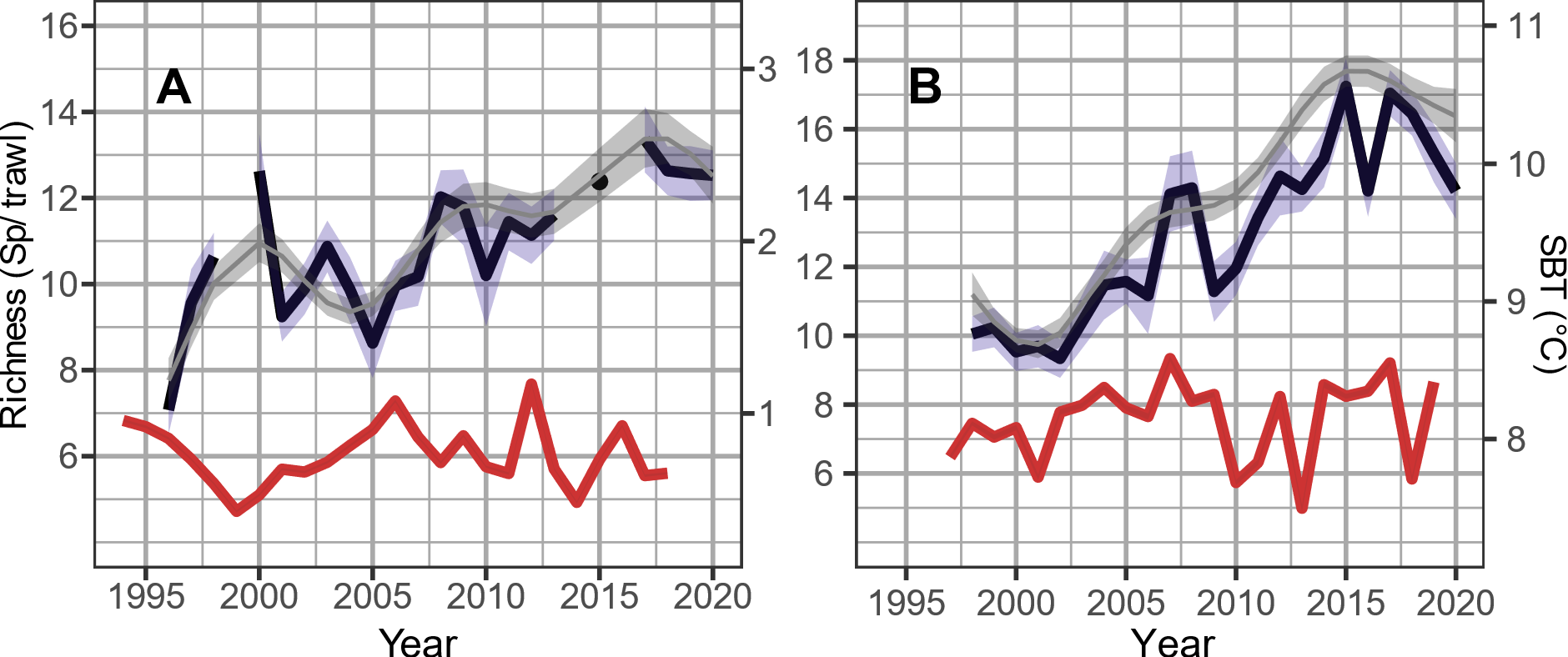

**Figure S1.** Average species per trawl (alpha diversity) across time in adjacent areas A: Around Svalbard; B: North Sea. Black line represents mean species richness per trawl with 95% CI in blue. Grey smoothed line and light grey 95 %CI represents the marginal effect of Year for constant sampling effort from a GAM model using Year and swept area as an offset (**Table 1**). Red line indicates changes in mean Sea Bottom Temperature across the whole area (correlation with SBT; p > 0.05).

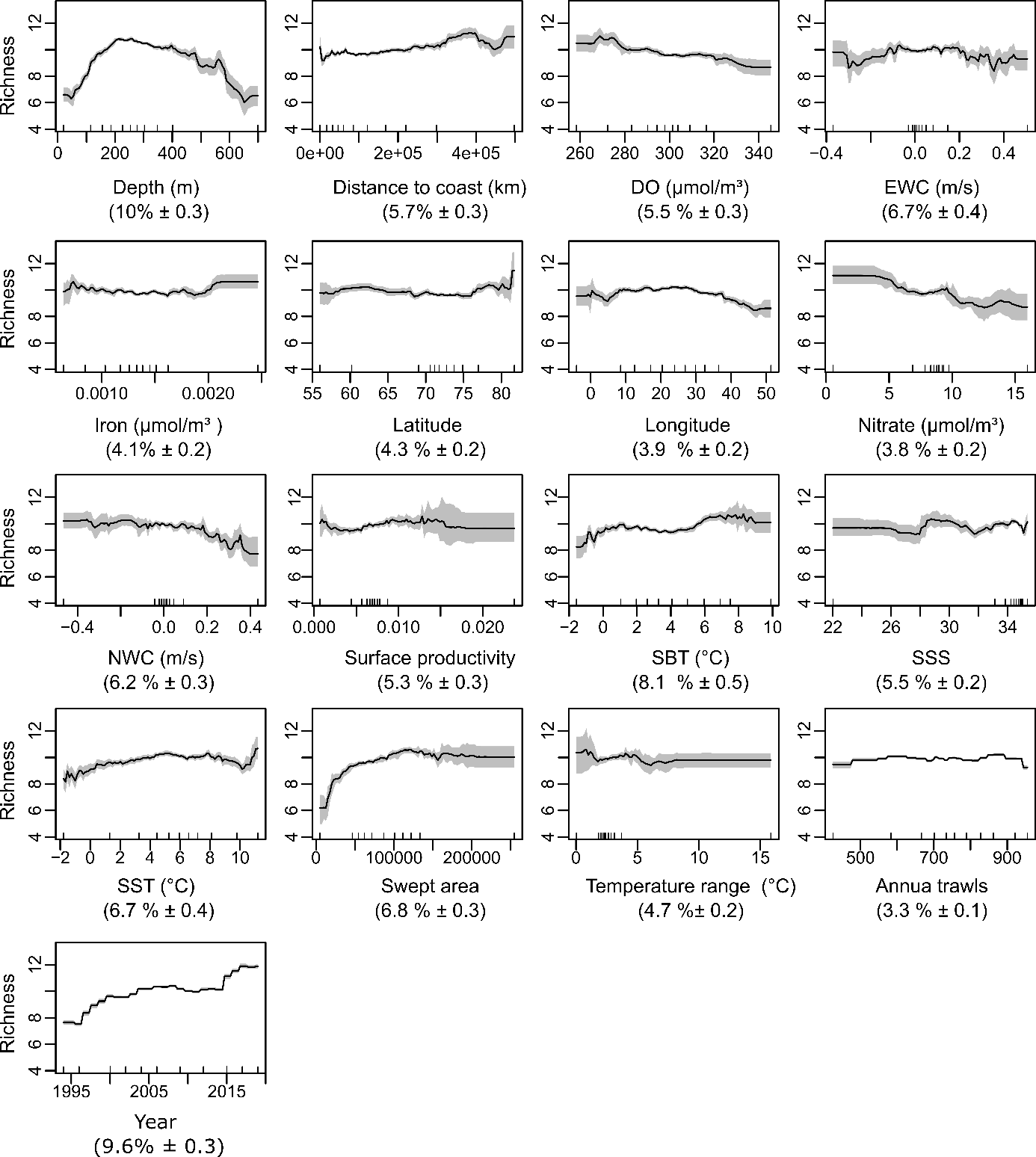

**Figure S2.** Partial dependence plots of the BRT model from 1994 to 2019. Between parenthesis is the relevance of each variable in the model.

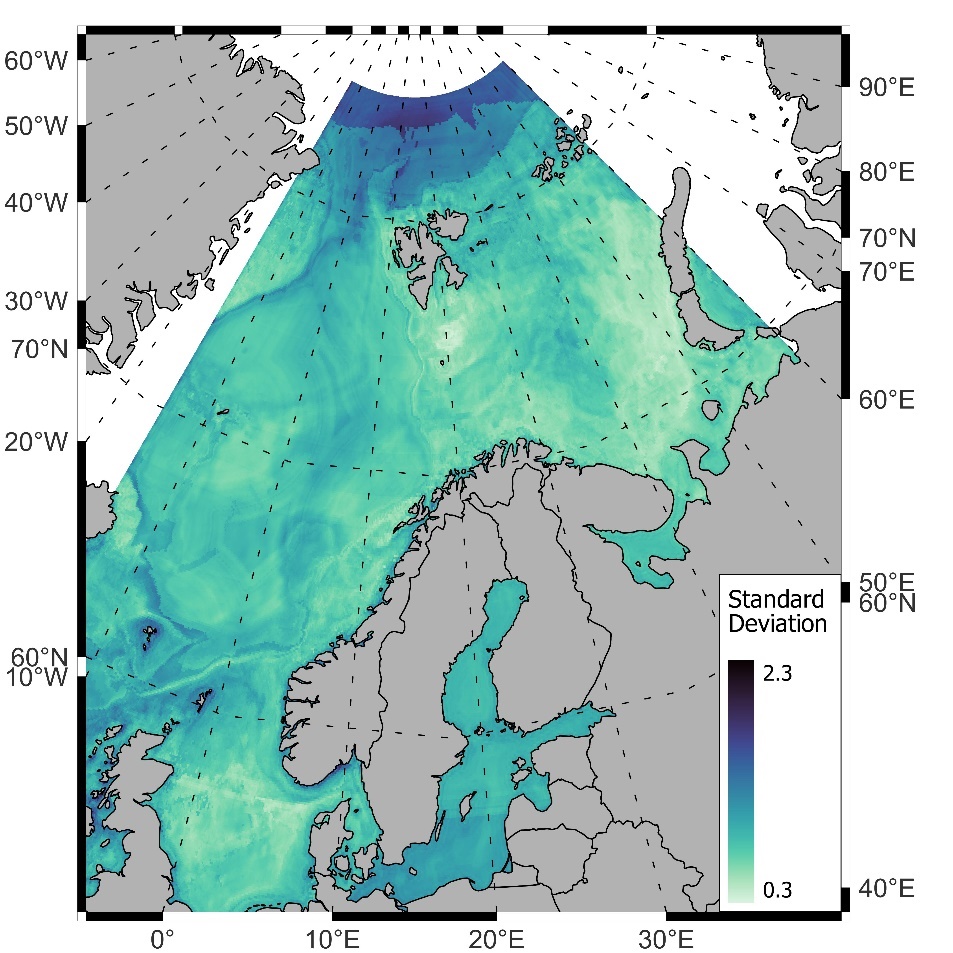

**Figure S3.** Overall mean standard deviation from BRT annual species richness predictions.

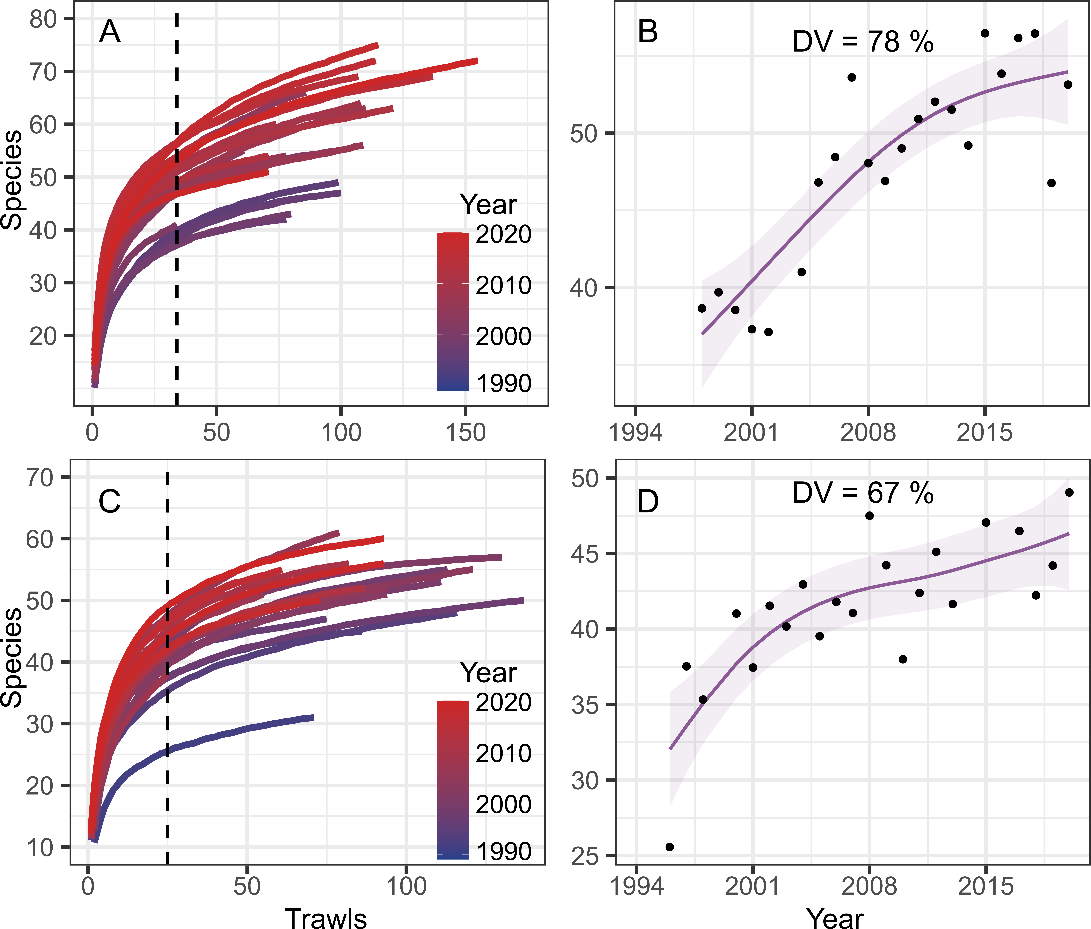

**Figure S4.** Gamma diversity across time in adjacent areas A: North Sea annual (rarefacted) SACs; B: North Sea species richness at minimum common number of sites (n = 34); C: Svalbard annual (rarefacted) SACs; D: Svalbard species richness at minimum common number of sites (n = 25).

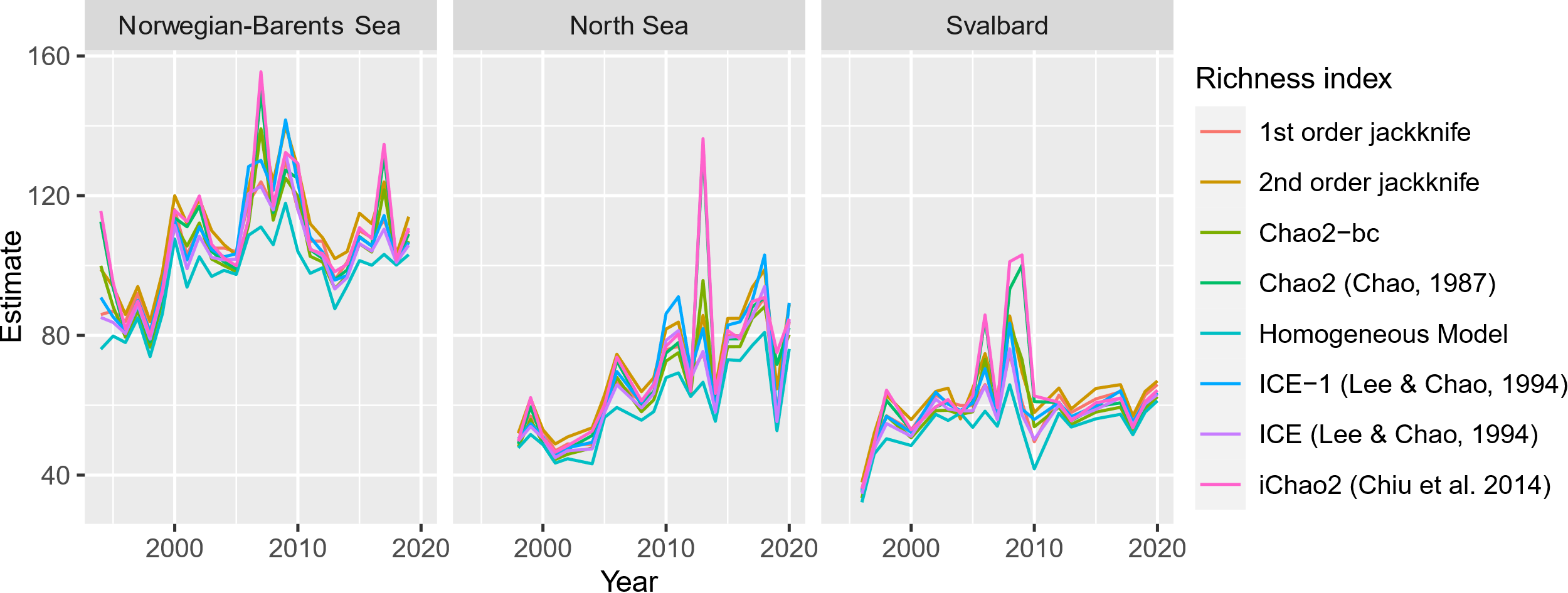

**Figure S5**. Chao indices change across study areas and years.

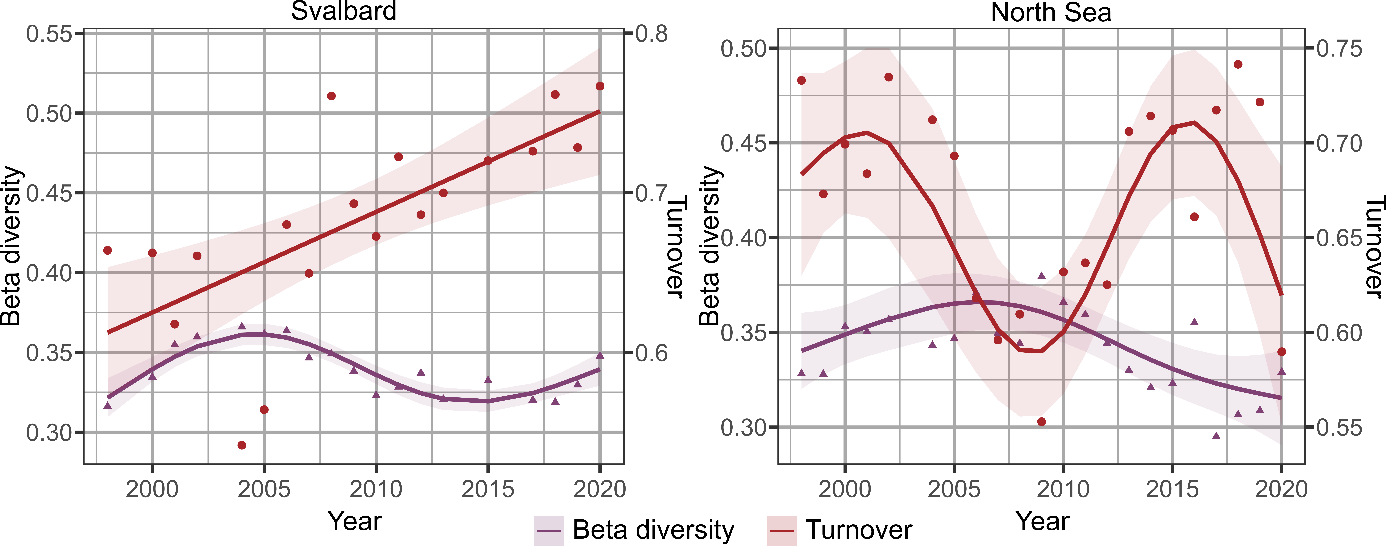

**Figure S6.** Annual change in beta diversity and turnover at adjacent regions, using mean pairwise Jaccard dissimilarity index.

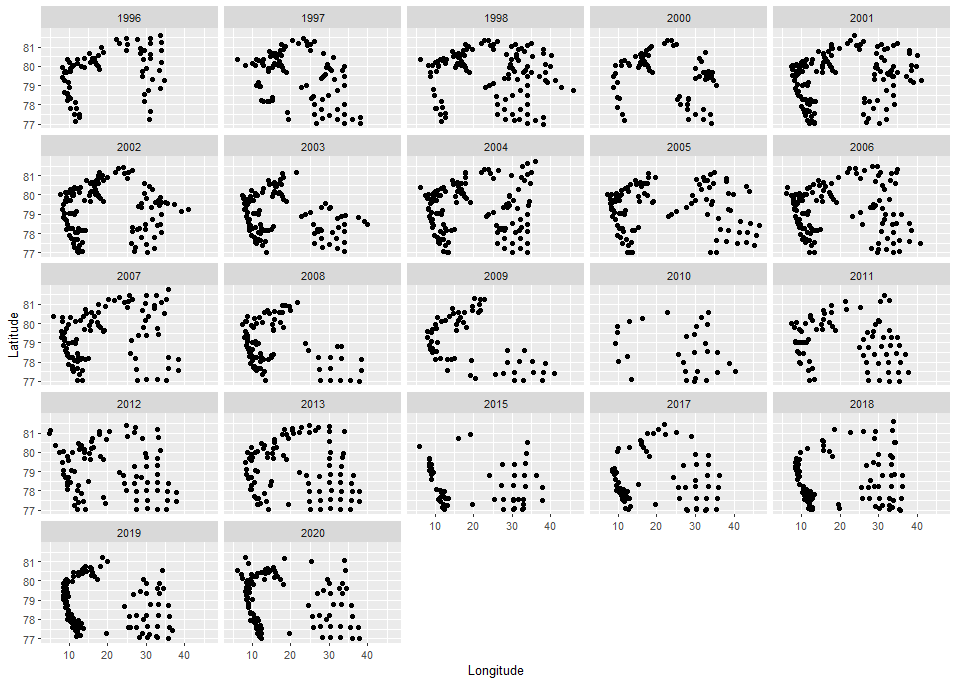
**(A)**

**(B)**

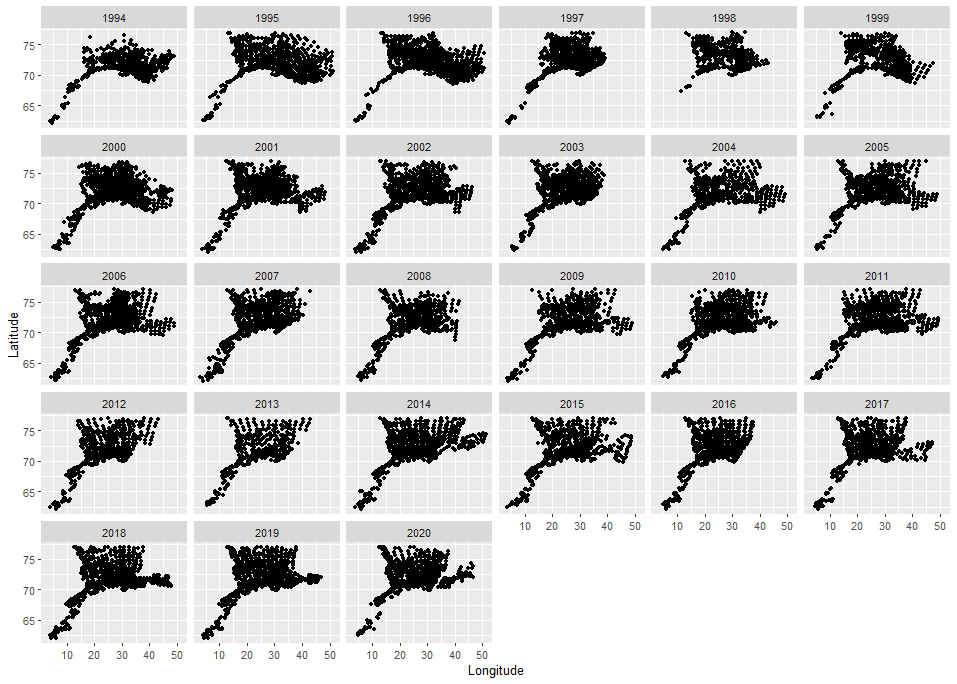

**(C)**

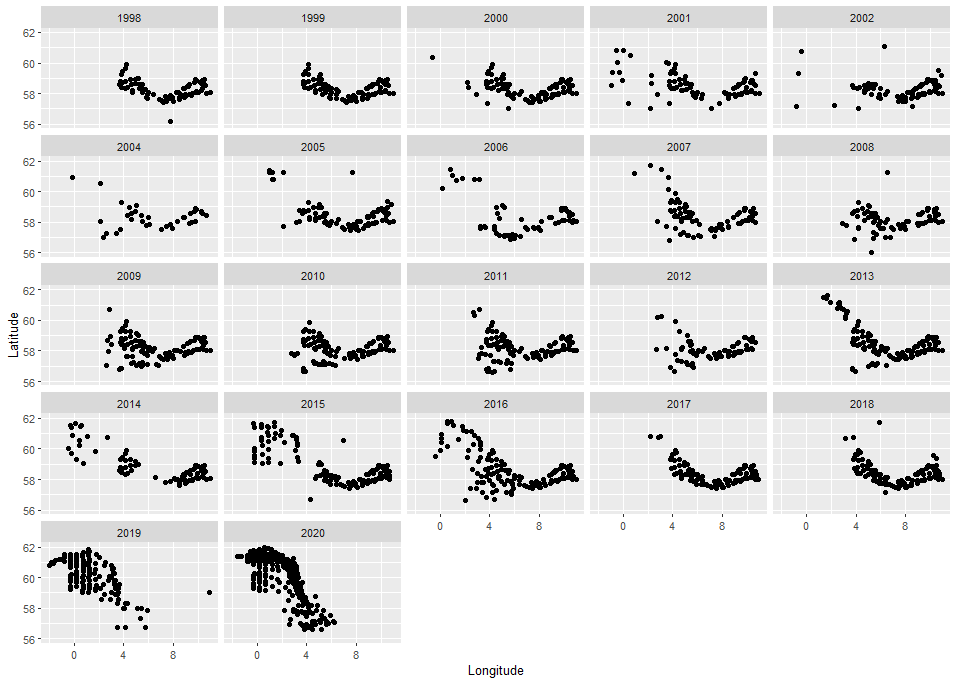

**Figure S7.** Spatial and temporal distribution of the sampling trawls. A: Main study area Norwegian-Barents Sea; B: Adjacent region around Svalbard; C: Adjacent region in the North Sea**.**

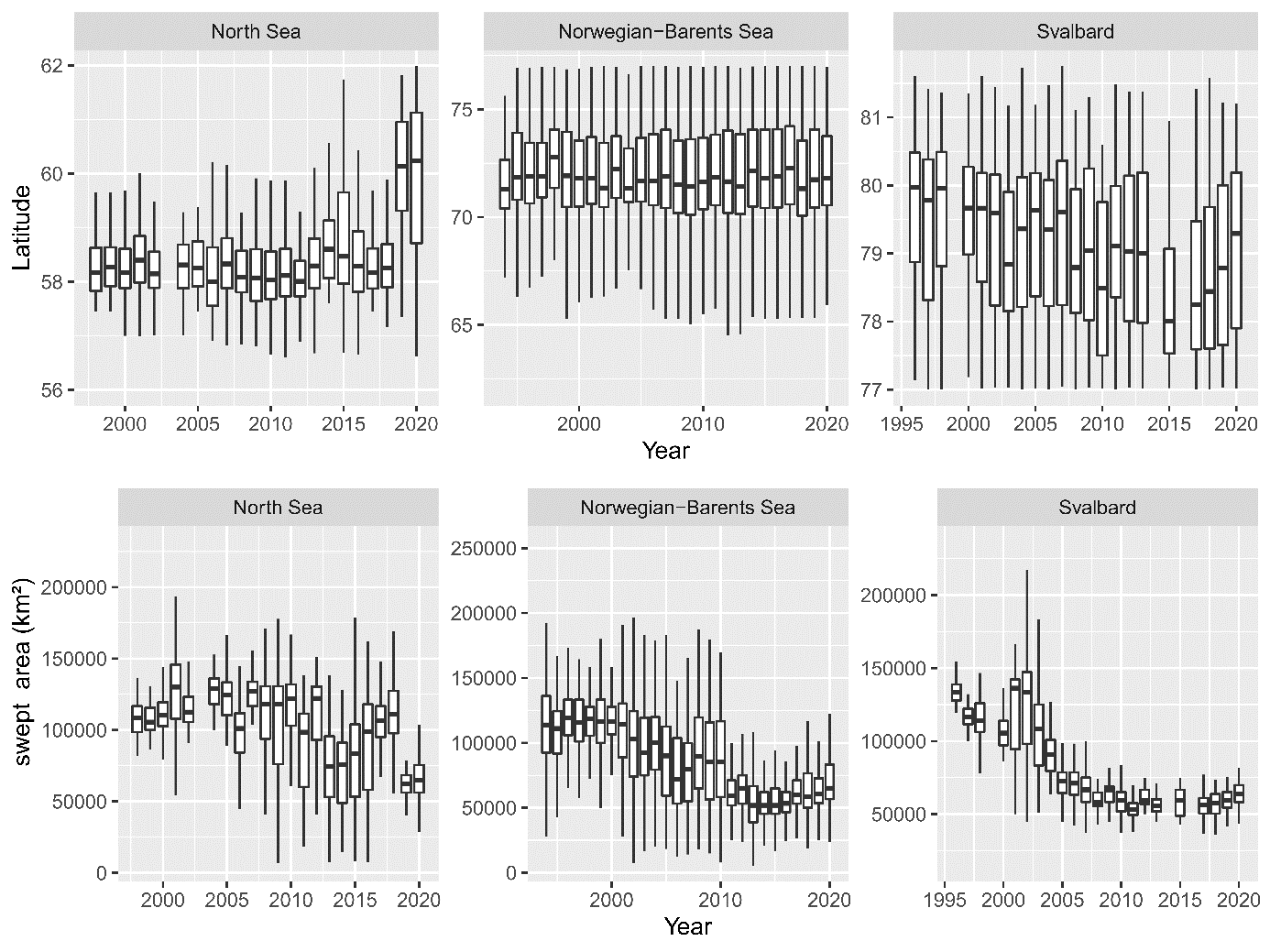

**Figure S8.** Dataset exploration graphics. Top: Latitudinal distribution of trawls, per year; Bottom: swept area (effort) at each trawl, per year.

**Table S1.** Existing studies of fish species richness changes in the Arctic Oceans.

| **Ref** | **Region** | **Low lat**  **(° N)** | **High lat (° N)** | **No. spp** | **Years** | **Comments** |
| --- | --- | --- | --- | --- | --- | --- |
| Batt et  al., 2017 | East Bering Sea (1984–2014, n = 31)  Aleutian Islands (1983–2014, n = 12),  Gulf of Alaska (1984–2013, n = 13),  West Coast US (1977–2004, n = 10),  Gulf of Mexico (1984–2000, n = 17),  Southeast US (1990–2014, n = 25),  Northeast US (1982–2013, n = 32),  Scotian Shelf (1970–2010, n = 41),  Newfoundland (1996–2011, n = 16) | 20 | 60 | 581 | 1970-2013 | No Arctic latitudes. Regional increases of 30. |
| Chaudhary et al., 2021 | Global |  |  | 50,000 globally | 1955- 2019 | Suggested poleward shift spp. in northern hemisphere but high variability in data due to diversity of datasets used. Species shifting not reported. |
| Burrows et al., 2019 | North Atlantic and Pacific | 20 | 62 | > 80 |  | Report general increase in community thermal index in the North Atlantic, though mixed effects at 2 degrees latitudinal cells. Community level, no taxonomic resolution. |
| Sunday et al., 2012 | Global | - | - | 142 marine and terrestrial | 1650 - 2011 | Shifts at both range boundaries  No marine shifts reported in polar latitudes. |
| Pinsky et al., 2013 | East Bering Sea (1982–2011),  Aleutian Islands (1980–2010),  Gulf of Alaska (1984–2011),  West Coast US (1977–2004),  Gulf of Mexico (1987–2011),  Southeast US (1971–2009),  Northeast US (1968–2008),  Scotian Shelf (1970–2011),  Newfoundland (1995–2011) | 27 | 60 | 360 (maximum 107 spp. per region) | 1968 - 2011 | Marine taxa distribution shifts follow local climate velocities instead of a general pattern. No reference to richness. |
| Poloczanska et al., 2013 | Global | - | - | 857 |  | No studies reporting fish distributional shifts in polar regions. |
| Fossheim et al., 2015 | Barents Sea | 68 | 81 | 74 | 2004, 2012 | Increase in species richness between two years. No comparison to temperature. |
| Kortsch et al., 2015 | Barents Sea | 68 | 81 | 233 | 2004, 2012 | Food-web structure analysis of changes reported in Fossheim et al. 2015. |
| Frainer et al., 2017 | Barents Sea | 68 | 81 | 52 | 2004, 2012 | Same dataset than Fossheim et al. 2015. Analysis of functional traits. |
| Frainer et al., 2021 | Barents Sea | 68 | 81 | 49 | 2004-2017 | They report an increase in Arctic Barents Sea and stable richness in boreal Barents Sea, though only 49 species selected for the analyses, limited to the species with known functional traits. |
| Husson et al., 2022 | Barents Sea | 68 | 81 | 33 | 2004-2017 | Increase in abundance and extent of most species during 2/3 extremely warm periods within this time period. Change in mean long and lat in the last 1/3. |
| Pecuchet et al., 2020 | Barents Sea | 68 | 81 | 239 | 2014-2017 | 11 boreal species observed in the Arctic region of the Barents Sea |
| Hiddink & ter Hofstede, 2008 | North Sea | 51 | 62 | 118 | 1985-2006 | Similar research trawl data to present study, and similar magnitude. |
| Fisher et al., 2008 | North-western pacific | 35 | 55 | 133 | 1973 -2003 | Correlation of -0.45 between richness and NAO. No long-term trend |
| Drinkwater & Kristiansen, 2018 | North Atlantic and Arctic | 35 | 81 | 12 fish species | 1950-2000 | Links changes in recruitment to AMO index and cold period during 1970s – 1980s |
| Tittensor et al., 2021 | Global | - | - | many taxa, depending on model used | 2000-2100 | Ensemble of 9 Marine Ecosystem Models. No species taxonomic resolution. Predict a decline in biomass in the Norwegian and boreal Barents Sea. Increase in biomass in the Arctic Barents. |
| Coll et al., 2020 | Global | - | - | ~ 3400 taxa | 1950-2100 | EcoOcean Marine Ecosystem Model with 3400 species organized in functional groups. Predict a decline in consumers’ biomass in the North Atlantic and increase in biomass in the Arctic Ocean using IPSL model forcings. |
| Cheung et al., 2013 | Global | - | - | 990 | 1970-2006 | No taxonomic resolution. Increase in mean temperature of the catch worldwide. |
| Cheung et al., 2009 | Global | - | - | 836 fish + 230 inverts | 2050 | Model projection of species invasion in the arctic region of the Barents Sea, and species extinction in the Norwegian Sea and south of Svalbard by 2050. |
| Skaret et al., 2015 | Norwegian Sea | 66 | 69 |  | 2013 | Mackerel north expansion. |
| Druon et al., 2016 | Mediterranean Sea, North Atlantic and Gulf of Mexico | 12 | 72 | 1 | 2003-2012 | Ecological niche modelling of Atlantic bluefin tuna. |
| Alabia et al., 2021 | Eastern Bering Sea | 55 | 62 | 159 (66 fish, 93 inverts) | 1990 - 2018 | Very similar trend than ours in Alpha diversity, but in markedly smaller latitudinal range, and not polar. |
| (Brunel & Boucher, 2007) | North-east Atlantic | 45 | 70 | 9 fish (4 species at high latitudes) | 1970-1998 | Most recruitment declining in the Baltic Sea, north Sea, west of Scotland and Irish populations. Herring and populations of boreal ecosystems followed an increasing trend. |
| Mueter & Litzow, 2008 | Berings Sea | 54 | 61 | 46 fish and inverts |  | Links community changes to sea ice retreat. Reports an increase in sp richness < 20 %. |
| Magurran et al., 2015 | North Sea | 55 | 59 | 131 (126 fish) | 1985-2013 | No change in species richness, though increase in turnover (beta diversity), with time. |
| Molinos et al., 2016 | Global | - | - | 13,000 marine species | 2006-2100 | Projected increase in richness (alpha diversity) across our study area until 2100. Unclear trend in dissimilarity (beta diversity). |
| Sunday et al., 2015 | Eastern Australia | 20 | 45 | 50 fish + 54 inverts | 1945-2005 | Poleward species shifts. |
| Morley et al., 2018 | Northeastern Pacific & Northwestern atlantic | 27 | 60 | 686 species | 2007-2100 | Mostly poleward shifts projected. Projections mostly based on the data from Batt et al. 2017 and Pinsky et al. 2013. |
| Pilotto et al., 2020 | North Atlantic | 47 | 61 | 6200 marine, freshwater and terrestrial |  | Increases in richness and abundance with increasing temperatures in most biogeoregions of Northern and Eastern Europe. |
| Pereira et al., 2010 | Global | - | - | 1066 fish and inverts | 2005-2050 | Rate of projected poleward shift among the demersal community are highest in the North Atlantic and Barents Sea. |

**Table S2.** Taxonomic classification of the species found.

| **Class** | **Order** | **Family** | **N especies** |
| --- | --- | --- | --- |
| Actinopteri | Acanthuriformes | Caproidae | 1 |
| Actinopteri | Anguilliformes | Anguillidae | 1 |
| Actinopteri | Anguilliformes | Congridae | 1 |
| Actinopteri | Argentiniformes | Argentinidae | 2 |
| Actinopteri | Argentiniformes | Microstomatidae | 1 |
| Actinopteri | Aulopiformes | Paralepididae | 2 |
| Actinopteri | Beloniformes | Belonidae | 1 |
| Actinopteri | Beloniformes | Scomberesocidae | 1 |
| Actinopteri | Callionymiformes | Callionymidae | 3 |
| Actinopteri | Carangiformes | Carangidae | 1 |
| Actinopteri | Clupeiformes | Clupeidae | 3 |
| Actinopteri | Clupeiformes | Engraulidae | 1 |
| Actinopteri | Eupercaria/misc | Moronidae | 1 |
| Actinopteri | Gadiformes | Gadidae | 13 |
| Actinopteri | Gadiformes | Gaidropsaridae | 6 |
| Actinopteri | Gadiformes | Lotidae | 3 |
| Actinopteri | Gadiformes | Macrouridae | 4 |
| Actinopteri | Gadiformes | Merlucciidae | 1 |
| Actinopteri | Gadiformes | Phycidae | 2 |
| Actinopteri | Gadiformes | Ranicipitidae | 1 |
| Actinopteri | Gobiiformes | Gobiidae | 4 |
| Actinopteri | Lophiiformes | Lophiidae | 1 |
| Actinopteri | Mulliformes | Mullidae | 1 |
| Actinopteri | Myctophiformes | Myctophidae | 3 |
| Actinopteri | Ophidiiformes | Carapidae | 1 |
| Actinopteri | Osmeriformes | Osmeridae | 1 |
| Actinopteri | Perciformes/Cottoidei | Agonidae | 3 |
| Actinopteri | Perciformes/Cottoidei | Cottidae | 11 |
| Actinopteri | Perciformes/Cottoidei | Cyclopteridae | 4 |
| Actinopteri | Perciformes/Cottoidei | Liparidae | 16 |
| Actinopteri | Perciformes/Cottoidei | Psychrolutidae | 2 |
| Actinopteri | Perciformes/Gasterosteoidei | Gasterosteidae | 1 |
| Actinopteri | Perciformes/Percoidei | Trachinidae | 2 |
| Actinopteri | Perciformes/Scorpaenoidei | Sebastidae | 4 |
| Actinopteri | Perciformes/Scorpaenoidei | Triglidae | 3 |
| Actinopteri | Perciformes/Uranoscopoidei | Ammodytidae | 4 |
| Actinopteri | Perciformes/Zoarcoidei | Anarhichadidae | 3 |
| Actinopteri | Perciformes/Zoarcoidei | Lumpenidae | 4 |
| Actinopteri | Perciformes/Zoarcoidei | Pholidae | 1 |
| Actinopteri | Perciformes/Zoarcoidei | Stichaeidae | 1 |
| Actinopteri | Perciformes/Zoarcoidei | Zoarcidae | 18 |
| Actinopteri | Pleuronectiformes | Bothidae | 1 |
| Actinopteri | Pleuronectiformes | Pleuronectidae | 8 |
| Actinopteri | Pleuronectiformes | Scophthalmidae | 6 |
| Actinopteri | Pleuronectiformes | Soleidae | 2 |
| Actinopteri | Salmoniformes | Salmonidae | 2 |
| Actinopteri | Scombriformes | Centrolophidae | 1 |
| Actinopteri | Scombriformes | Scombridae | 1 |
| Actinopteri | Stomiiformes | Sternoptychidae | 2 |
| Actinopteri | Syngnathiformes | Syngnathidae | 2 |
| Actinopteri | Zeiformes | Zeidae | 1 |
| Elasmobranchii | Carcharhiniformes | Pentanchidae | 1 |
| Elasmobranchii | Carcharhiniformes | Scyliorhinidae | 2 |
| Elasmobranchii | Carcharhiniformes | Triakidae | 1 |
| Elasmobranchii | Rajiformes | Arhynchobatidae | 1 |
| Elasmobranchii | Rajiformes | Rajidae | 13 |
| Elasmobranchii | Squaliformes | Centrophoridae | 1 |
| Elasmobranchii | Squaliformes | Etmopteridae | 1 |
| Elasmobranchii | Squaliformes | Somniosidae | 1 |
| Elasmobranchii | Squaliformes | Squalidae | 1 |
| Holocephali | Chimaeriformes | Chimaeridae | 1 |
| Myxini | Myxiniformes | Myxinidae | 1 |
| Petromyzonti | Petromyzontiformes | Petromyzontidae | 1 |
|  |  |  | 188 |

**Table S3.** Variables used in the BRT model. Variables with a (*) were excluded after proving not relevant in the exploration partial dependence plots. (< 3 % of DV explained).

| **Layer name** | **Layer code** | **Units** | **Origin** | **Temporal resolution** |
| --- | --- | --- | --- | --- |
| Sea surface temperature | SST | °C | Copernicus | Yes |
| Sea surface salinity | SSS | psu | Copernicus | Yes |
| Sea bottom temperature | SBT | °C | Copernicus | Yes |
| Northward currents | NWC | m/s | Copernicus | Yes |
| Eastward currents | EWC | m/s | Copernicus | Yes |
| Ice concentration * | ICC | Area fraction 0 to 1 | Copernicus | Yes |
| Bottom nitrate concentration | Nitrate | μmol/m^3^ | Bio Oracle | No |
| Bottom oxygen concentration | DO | μmol/m^3^ | Bio Oracle | No |
| Bottom iron concentration | Iron | μmol/m^3^ | Bio Oracle | No |
| Surface productivity | Prod.Surface | g/m^3^/day | Bio Oracle | No |
| Temperature Range | Temp. range | °C | Bio Oracle | No |
| Bottom productivity * | Bot. Prod | g/m^3^/day | Bio Oracle | No |
| Bathymetry | Depth | M | MARSPEC | No |
| Distance to coast | Dist_to_coast | M | Self made | No |
| Longitude | Longitude | ° | Self made | No |
| Latitude | Latitude | ° | Self made | No |
| Year | Year | yr | Self made | Yes |
| Swept area | swept_area | Km^2^ | Self made | No |
| Annual trawl | Annual_trawls | Trawls | Self made | No |
